# Effective connectivity of the human claustrum: Triple networks, subcortical circuits, and psychedelic modulation

**DOI:** 10.1101/2025.09.07.674759

**Authors:** Navid Shams Masjedi, Adeel Razi

## Abstract

Decades of cross-species research highlight the claustrums extensive bidirectional connectivity with cortical and subcortical regions, implicating it in higher-order cognitive processes requiring synchronized brain states. Psychedelics may disrupt this synchrony by modulating claustro-cortical signaling, reflected by the dissolution of cortical network signatures. Using spectral dynamic causal modeling on resting-state fMRI data from the Human Connectome Project and PsiConnect datasets at 7T and 3T, we provide the first in vivo characterization of claustral effective connectivity with triple networks and subcortical regions in humans, both at rest and under the influence of psilocybin. Claustra displayed widespread bidirectional effective connectivity and a strong inhibitory influence on all target regions. Psilocybin enhanced claustral inhibition of cortical networks while disinhibiting subcortical areas, partially associated with psychedelic subjective effect scores. These findings are consistent with cellular and functional cross-species data, supporting the proposed mechanism of claustro-cortical inhibition in regulating network synchrony, while extending this influence to the subcortex, and revealing hierarchical and hemispheric asymmetries in claustral signaling modulation under psilocybin.

## Introduction

The claustrum (CLA) remains one of the brain’s most enigmatic structures, with its functional role in health and disease still unresolved (Benarroch, 2021; Evleksiz et al., 2025). It is a bilateral telencephalic nucleus conserved across species, forming a thin, curved sheet of gray matter situated between the putamen and insula (see Fig. 1**a**) ((Bruguier et al., 2020; Kowianśki et al., 1998; Smith et al., 2020)). Since its landmark proposal as a neural correlate of consciousness (NCC) (Crick and Koch, 2005), the CLA has been implicated in diverse higher-order functions, from attentional salience (Atlan et al., 2018; Goll et al., 2015; Remedios et al., 2014; Smith et al., 2019; Smythies et al., 2014), to sensorimotor processing (Chevée et al., 2022; Jankowski and O’Mara, 2015; Ollerenshaw et al., 2021; Smith and Alloway, 2014; Smith et al., 2012), learning and memory (Bhattacharjee et al., 2024; Medina et al., 2024), and multiple dimensions of consciousness (Liaw and Augustine, 2023), including anaesthesia (Bosinski and Connor, 2025; Luo et al., 2023; Pavel et al., 2019; Smith et al., 2017), sleep (Atlan et al., 2024; Lamsam et al., 2024; Narikiyo et al., 2020; Norimoto et al., 2020), and pharmacologically altered states (Doss et al., 2022; Stiefel et al., 2014). Similarly, the CLA has received increased attention as a clinical target, with morphological and functional alterations reported in epilepsy (Kurada et al., 2019; Steriade et al., 2017; Watson et al., 2024), Alzheimer’s disease (Ayyildiz et al., 2023; Chen et al., 2023; Venneri and Shanks, 2014), Parkinson’s disease (Arrigo et al., 2019; Kalaitzakis et al., 2009; Sitte et al., 2017; Zedde et al., 2025), schizophrenia (Cascella et al., 2011; Cascella and Sawa, 2014; Galeno et al., 2004; Patru and Reser, 2015; Schinz et al., 2024), and chronic pain (Stewart et al., 2024; Zhang and Zamponi, 2024).

**Fig. 1:**
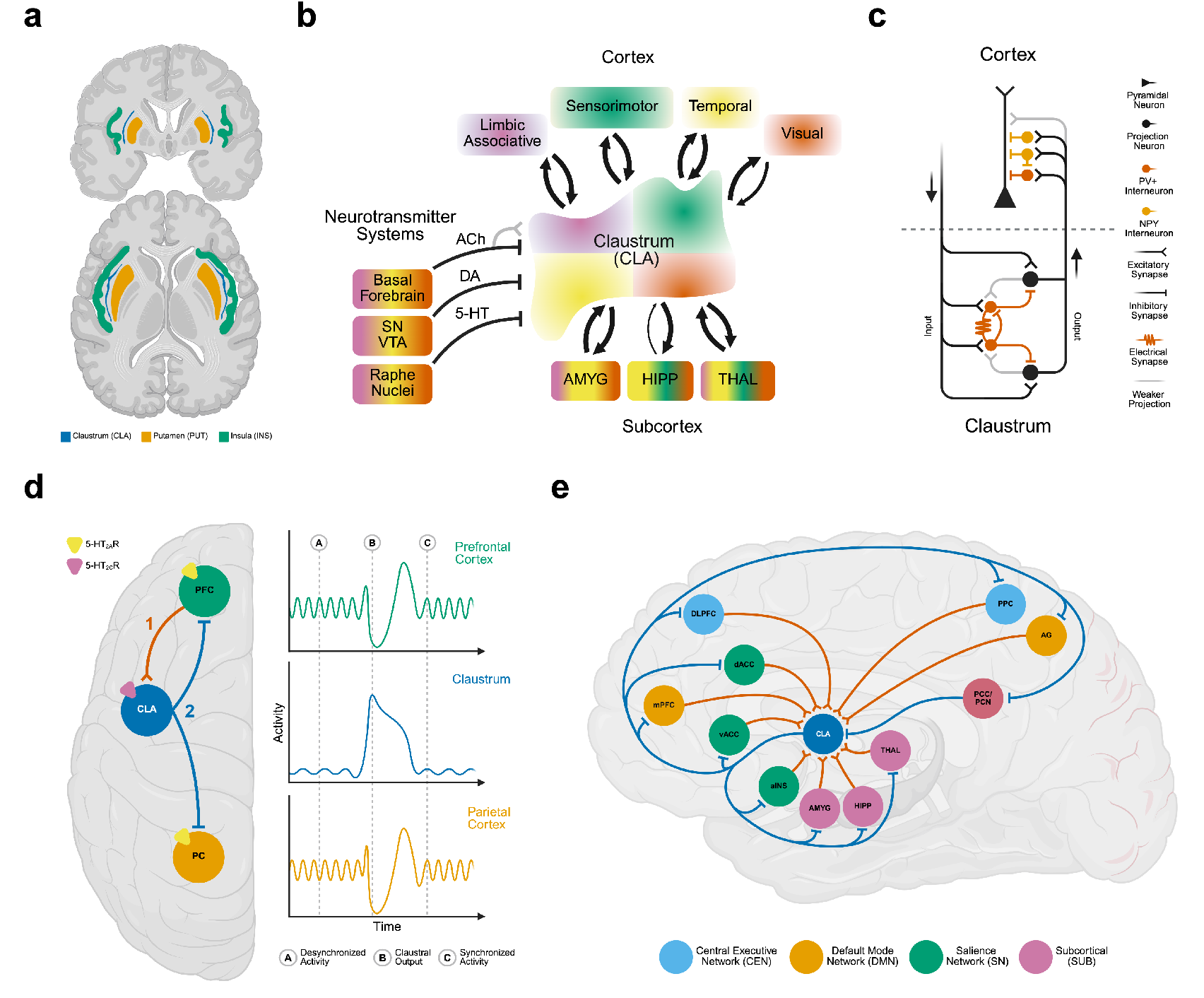
Present understanding of claustrum (CLA) connectivity and function. **a** Anatomical localization of the CLA. **b** Structural connectivity: Animal studies reveal extensive bidirectional projections between CLA, cortex, and subcortex following a topographical organization. The CLA also receives input from multiple neurotransmitter systems, largely inhibitory. **c** Cellular connectivity: Cortical afferents target excitatory projection neurons and abundant parvalbumin (PV) interneurons. PV cells are densely interconnected via gap junctions, forming feedforward inhibition (FFI) microcircuits onto CLA efferents. CLA projection neurons strongly innervate cortical neuropeptide Y (NPY) and PV interneurons, and to a lesser extent pyramidal cells. Thus, cortico-claustral input can excite or inhibit the CLA, while claustro-cortical output primarily induces pyramidal FFI. **d** Cortical influence: Functional hypotheses suggest that frontal cortices direct the CLA to instantiate cortical network states via concurrent inhibition of prefrontal and parietal nodes. This way, the CLA couples its high-frequency activity to lower cortical rhythms, phase-locking distal oscillations. Psychedelics are hypothesized to disrupt this function, dissolving network signatures via 5-HT_2A/C_R-mediated mechanisms. **e** Human effective connectivity: Our results replicate animal findings, showing widespread bidirectional CLA–cortex and CLA–subcortex influences. The CLA inhibited all cortical and subcortical targets while receiving mostly excitatory input, with exception of the posterior parietal cortex/precuneus (PCC/PCN) region, which uniquely inhibited the CLA.

Current functional theories bridge these widespread implications by positing the CLA as a key hub for the coordination of cortical networks and synchronized brain states (Do et al., 2024; Madden et al., 2022; Vidyasagar and Levichkina, 2019). The hallmark feature of the CLA is its widespread bidirectional structural connectivity to almost the entire cortex across species (Jackson et al., 2020), more recently extended to subcortical structures such as the amygdala, hippocampus, and thalamus (Lei et al., 2025). Furthermore, the CLA receives projections from various endogenous neurotransmitter systems, primarily inhibiting its activity (see Fig. 1**b**) (Wong et al., 2021).

Cellularly, the CLA follows a core–shell structure that is topographically organized and divided into functional modules (Chia et al., 2020; Erwin et al., 2021; Lei et al., 2025; Marriott et al., 2021; Zingg et al., 2018). It is primarily composed of glutamatergic projection neurons while housing a core of GABAergic interneurons expressing different calcium-binding proteins and peptides (Chong and Gămănuţ, 2024; Graf et al., 2020; Takahashi et al., 2023). Within the CLA, fast-spiking parvalbumin (PV) interneurons form highly interconnected feedforward inhibition (FFI) microcircuits reciprocally mapping onto cortical afferents and efferents (Graf et al., 2020; Kim et al., 2016). Furthermore, while CLA influence on the cortex is layer- and cell-type–specific (McBride et al., 2023; Wang et al., 2023), CLA efferents display a bias towards cortical neuropeptide Y (NPY) and PV interneurons, inducing FFI of pyramidal neurons (see Fig. 1**c**) (de la Torre-Martínez et al., 2025; Jackson et al., 2018).

In humans, anatomical and diffusion imaging confirm the CLA’s widespread projection system (Arrigo et al., 2017; Fernández-Miranda et al., 2008; Milardi et al., 2015; Wendt et al., 2024)), terming it the most densely interconnected region per unit volume in the human brain and a key contributor to cortical network architecture (Torgerson et al., 2015). CLA connectivity is especially pronounced to frontal cortices, which have been linked to cortical network motifs (Qadir et al., 2022; White and Mathur, 2018; White et al., 2018; Zahacy et al., 2024). Furthermore, the CLA displays significant functional connectivity (FC) to resting-state networks, including the frontoparietal central executive (CEN), default mode (DMN), and salience network (SN) (Holzscherer et al., 2025; Krimmel et al., 2019; Rodríguez-Vidal et al., 2024). More importantly, CLA activation has been shown to coincide with activation of task-positive network signatures across cognitively demanding tasks (Huang et al., 2024; Krimmel et al., 2019). Thus, the CLA has been implicated in a cognitive control framework, termed the network instantiation in cognitive control (NICC) model, which posits that frontal cortices direct the CLA to instantiate cortical network signatures by synchronizing frontal–parietal activity via acute inhibition (see Fig. 1**d**) (Madden et al., 2022).

Based on this framework, the cortico-claustro-cortical (CCC) model of psychedelic action proposes modulation of this mechanism via serotonin 2A receptor (5-HT_2A_R)–mediated mechanisms (Doss et al., 2022). Psychedelic compounds, such as psilocybin and lysergic acid diethylamide (LSD), exert their action at a wide range of receptors (Jain et al., 2025), with 5-HT receptor subtypes being most strongly implicated in psychedelic subjective effects (Cameron et al., 2023; Erkizia-Santamaría et al., 2022). Converging evidence suggests that psychedelics disrupt cortical synchrony, manifesting in the dissolution of cortical network signatures (Shinozuka et al., 2024; Siegel et al., 2024; Yu et al., 2024), which has been proposed as a CLA-mediated therapeutic mechanism (Nichols et al.,2017). Initial findings in rodents identified a functionally distinct subpopulation of 5-HT_2A_R-expressing glutamatergic CLA neurons, which directly depolarized following administration of 2,5-dimethoxy-4-iodoamphetamine (DOI) (Martin and Nichols, 2016). More recent evidence in rodents suggests a preferential role for 5-HT_2C_Rs exclusively facilitating the induction of glutamate receptor–mediated excitatory postsynaptic currents (EPSCs) by DOI (Anderson et al., 2024). In humans, psilocybin has been shown to differentially modulate FC between the CLA and multiple resting-state networks asymmetrically, which was associated with measures of network integrity and modularity (Barrett et al., 2020). Furthermore, a head-to-head comparison of psilocybin and the highly selective kappa opioid receptor (KOR) agonist salvinorin A revealed convergent effects on the CLA, increasing FC to frontal cortices, while dissociating this shared action from changes in thalamo–cortical FC exclusive to psilocybin (Bagdasarian et al., 2024). Thus, the CLA may present a point of convergence for the action of classical psychedelics and the atypical hallucinogen salvinorin A (Stiefel et al., 2014), which induces changes in differential measures of FC strikingly similar to those by classical psychedelics in humans (Doss et al., 2020), and shares phenomenological aspects with psychedelic experiences (Brito-da Costa et al., 2021).

Taken together, the anatomical and cellular mapping of the CLA to the cortex, and the mechanism of claustro-cortical inhibition have been integral to current functional theories of claustral function (Doss et al., 2020; Madden et al., 2022; Vidyasagar and Levichkina, 2019). However, characterization of claustrum effective connectivity to major cortical networks and its input-output organization in humans has not been undertaken thus far. Additionally, its directed projection system to subcortical structures also remains to be characterized in humans, with divergent patterns reported in rodents and primates (Jackson et al., 2020; Lei et al., 2025). Furthermore, changes in effective connectivity between the CLA, cortex, and subcortex under psychedelic action have not been investigated yet.

Here, we employed spectral Dynamic Causal modeling (spDCM) on resting-state functional Magnetic Resonance Imaging (rs-fMRI) from the Human Connectome Project (HCP) at 3 and 7 Tesla (T) resolution, as well as the PsiConnect 3T (PC3T) dataset, to estimate the effective connectivity (EC) of the CLA to the Central Executive (CEN), Default Mode (DMN) and Salience (SN) networks, and subcortical regions, including the amygdala (AMYG), hippocampus (HIPP) and thalamus (THAL) (see Fig. 1**e**. Furthermore, we compared EC estimates from the baseline and administration sessions of the PC3T dataset to determine changes in EC under psilocybin, and their association with self-report measure scores of psychedelic subjective effects from the 30-item revised Mystical Experience Questionnaire (MEQ30) and 11-Dimension Altered States of Consciousness Questionnaire (11D-ASC) (see Fig. 2).

**Fig. 2:**
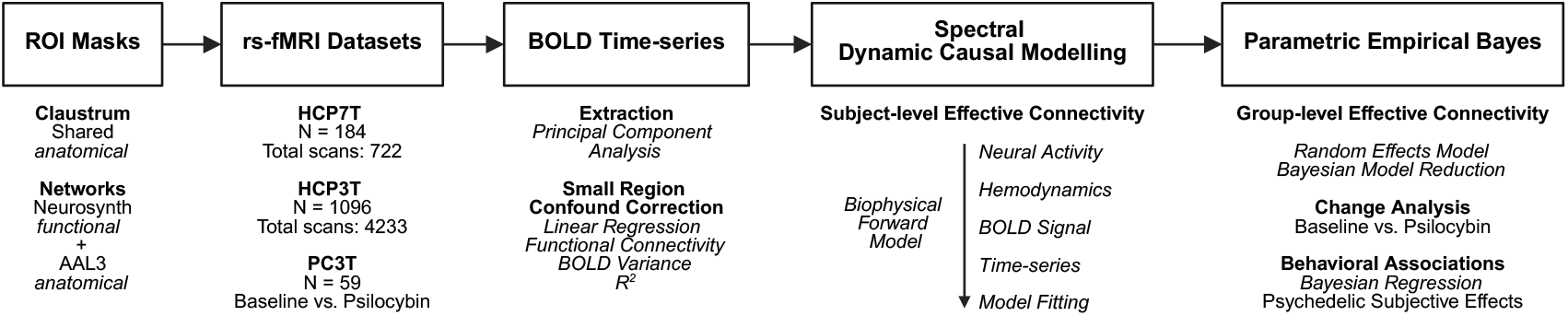
Analysis pipeline. Network masks were generated using functional activation maps constrained by anatomical atlas masks. Blood oxygen level-dependent (BOLD) time-series were extracted for all regions of interest (ROIs) across the three datasets, with small region confound correction (SRCC) applied to the CLA time-series. Spectral dynamic causal modeling (spDCM) estimated effective connectivity (EC) at the single-subject level for all networks, followed by group-level estimation via parametric empirical Bayes (PEB). For the Human Connectome Data (HCP) datasets, only mean EC was estimated. In PsiConnect (PC3T), change analyses were conducted comparing group-level baseline EC with EC under psilocybin. Additionally, connections displaying a change under psilocybin were tested for associations with psychedelic subjective effect scores using Bayesian regression.

## Results

### The claustrum exhibits widespread inhibition of cortical and subcortical regions

Across all datasets, DCM revealed a robust inhibitory influence of the bilateral CLA on all cortical and subcortical targets examined (see Fig. 3**a**). Cortical network inhibition strengths were largely comparable between datasets, with only minor cross-dataset variability observed for specific nodes, notably for weaker left CLA estimates of the CEN and SN in PC3T. Subcortical inhibition varied more substantially between datasets, still retaining estimates for all targets, which were consistently weaker than cortical inhibition within datasets.

**Fig. 3:**
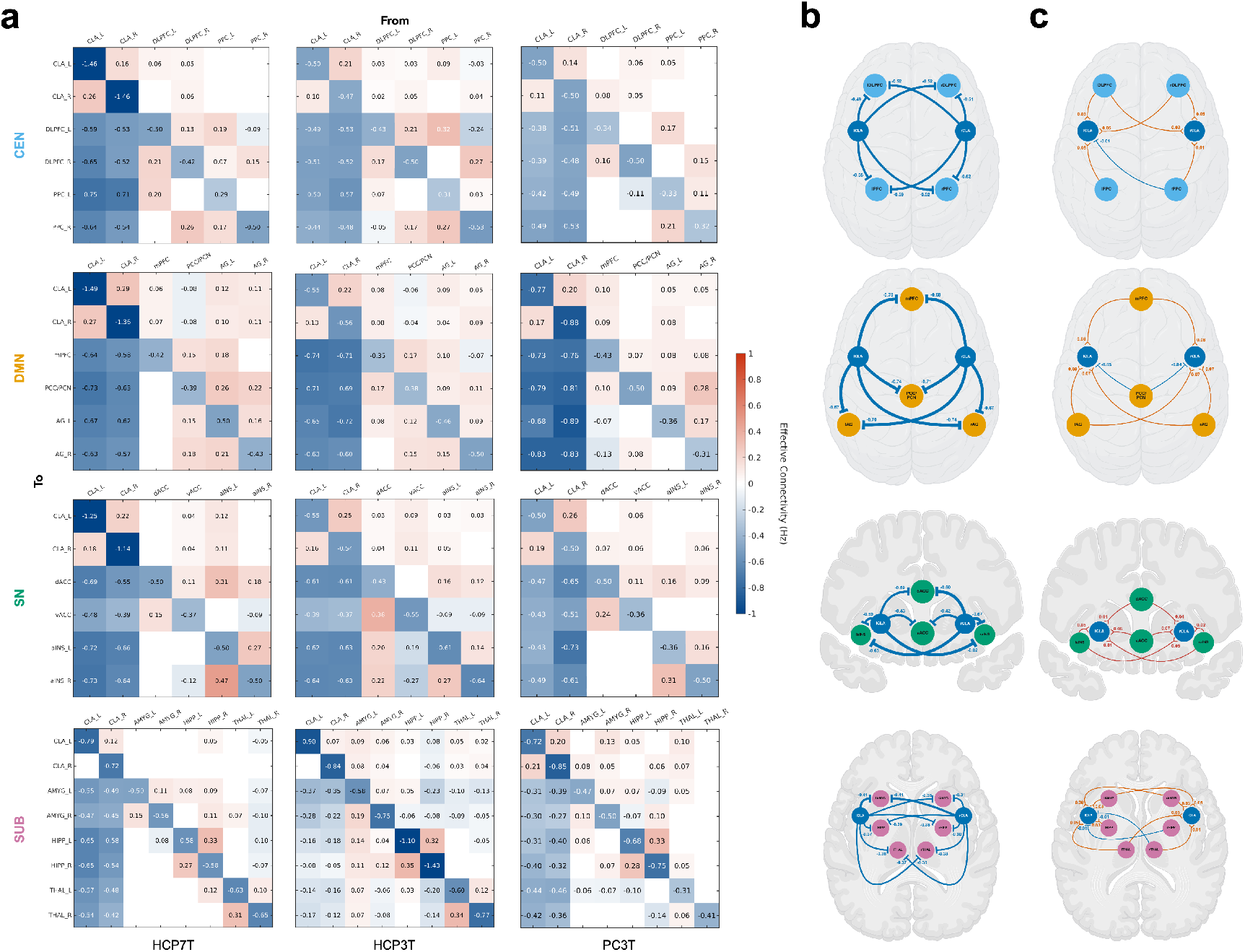
Effective connectivity (EC) at rest. **a** EC matrices across all networks and datasets. From row three onward, columns one and two show CLA outputs; from column three onward, rows one and two show inputs to the CLA. A schematic of the matix layout is provided in Fig. 1**a** for clarity. Values exceeding 50% of the matrix maximum are highlighted in white font. **b** Mean CLA output estimates per network. EC strength is reflected by the arrow thickness in incremenents of 0.20 Hz. **c** Mean CLA input estimates per network.

To evaluate overall output estimates of the CLA, we computed the mean EC across datasets (see Fig. 3**b**). Within cortical networks, CLA hemispheres displayed largely uniform inhibition strengths across targets. Only in the SN, the right CLA showed a minimally stronger influence on the ipsilateral aINS than the left CLA. Additionally, the vACC was less inhibited by both claustra than all other nodes. Between cortical networks, inhibition estimates were consistently stronger for the DMN (mean = -0.70 Hz), followed by the SN (mean = -0.57 Hz) and the CEN (mean = -0.53 Hz). Subcortical inhibition was overall weaker than cortical inhibition (mean = -0.36 Hz). Inhibition strengths per subcortical nodes were balanced across claustral hemispheres. The left AMYG was slightly stronger inhibited by both claustra, which, however, was due to the gradient in CLA inhibition observed in the HCP3T dataset, where the left AMYG was substantially stronger inhibited than all other subcortical ROIs (see Fig. 3**a**).

Overall, these findings indicate that the CLA exerts a uniformly inhibitory influence across the brain, with cortical targets being more strongly affected than subcortical ones, only subtle differences in hemispheric or regional specificity and slightly more pronounced DMN inhibition at rest.

### The claustrum receives diverse input from cortical and subcortical regions

CLA input estimates varied more substantially, with estimates differentially present between datasets across datasets and networks (see Fig. 3**a**). Overall, input estimates were much weaker than CLA output estimates and were predominantly excitatory. Inhibitory influences were estimated from the PCC/PCN across HCP datasets, from the right PPC in HCP3T, and divergent estimates were observed for the right THAL and right HIPP across datasets.

Mean input estimates per network across datasets reflect this more nuanced CLA input structure across networks (see Fig. 3**c**). Fully connected inputs were only estimated for the DMN and SN. SN nodes displayed exclusively excitatory inputs to both claustra. For the DMN, only the PCC/PCN displayed a mean inhibitory influence. The CEN displayed largely excitatory influences on both claustra. Notably, no input from the left PPC to the right CLA was estimated in any dataset. Similarly, the inhibitory influence of the right PPC on the left CLA was only present in HCP3T, with no estimates in the other two datasets. Subcortical regions also showed predominantly excitatory input to both claustra. Notably, the divergent estimates in excitatory and inhibitory valence for the right HIPP and THAL resulted in weak inhibitory influences on the contralateral CLA. Similarly, the influence from the right HIPP to the right CLA was abolished by equally strong divergent estimates in HCP3T and PC3T. No influence from the left HIPP to the left CLA was estimated in any dataset. Given the overall weak mean estimates for CLA input across cortex and subcortex, no hemispheric bias became apparent for any node. Likewise, input strengths were comparably strong across networks, with only marginally higher estimates for the DMN.

Thus, the CLA appears to integrate a balanced and predominantly excitatory set of generally weaker cortical and subcortical inputs at rest, with only the PCC/PCN region displaying a more consistent inhibitory influence over both claustra.

### Claustra display excitatory cross-hemispheric coupling and strong self-inhibition

Consistent bidirectional excitatory cross-claustral coupling was estimated across all datasets (see Fig. 3**a**). Cross-hemispheric coupling generally displayed comparable connection strengths within datasets. For the subcortical models of the HCP datasets, no left-to-right coupling was estimated. Mean cross-claustral coupling across models and datasets suggests a slightly stronger right-to-left influence (see Fig. 1**b**). CLA self-inhibition estimates showed a more varied pattern across datasets (see Fig. 3**a**). In HCP7T, CLA self-inhibition was exceptionally strong, often representing the largest connectivity estimate within networks. In HCP3T, CLA self-inhibition was only pronounced in the SUB model. In PC3T, relatively pronounced CLA self-inhibition was estimated in the DMN and SUB model. Mean CLA self-inhibition across models and datasets reveals overall strong self-inhibition of both claustra, at the same strength (see Fig. 1**b**).

Thus, we identified excitatory cross-hemispheric coupling between claustra, with a potentially slight bias from the right to left CLA, whereas both claustra display comparably strong self-inhibition,

### Psilocybin drives claustro-cortical inhibition asymmetrically

Using the administration session from the PC3T dataset as a covariate in the PEB framework, we compared EC across networks between the baseline and administration scan, identifying changes in CLA EC under psilocybin. Notably, psilocybin profoundly altered claustro-cortical dynamics, with the most prominent effect being an asymmetric upregulation of left CLA inhibition across cortical networks (see Fig. 4**a**). This effect was strongest for the CEN, with increased inhibition of the bilateral DLPFC and right PPC from the left CLA. Inhibition of contralateral targets was more pronounced than for the left DLPFC. No change in right CLA signaling to CEN nodes was detected. For the DMN, left CLA inhibition of the mPFC and left AG increased at comparable strengths, accompanied by a reduction in left-to-right CLA excitation. Contrarily, the right CLA disinhibited the right AG, also at a comparable magnitude, coinciding with slightly decreased right CLA self-inhibition. For the SN, the left CLA increased its inhibition of all nodes except the right aINS. This was especially pronounced for the left aINS, displaying the strongest increase in claustral inhibition across networks. Contrasting the other two networks, the right CLA also displayed increased inhibition of the dACC and left aINS, again slightly more pronounced for the latter. Notably, increased claustral inhibition presented the strongest change estimates across networks.

**Fig. 4:**
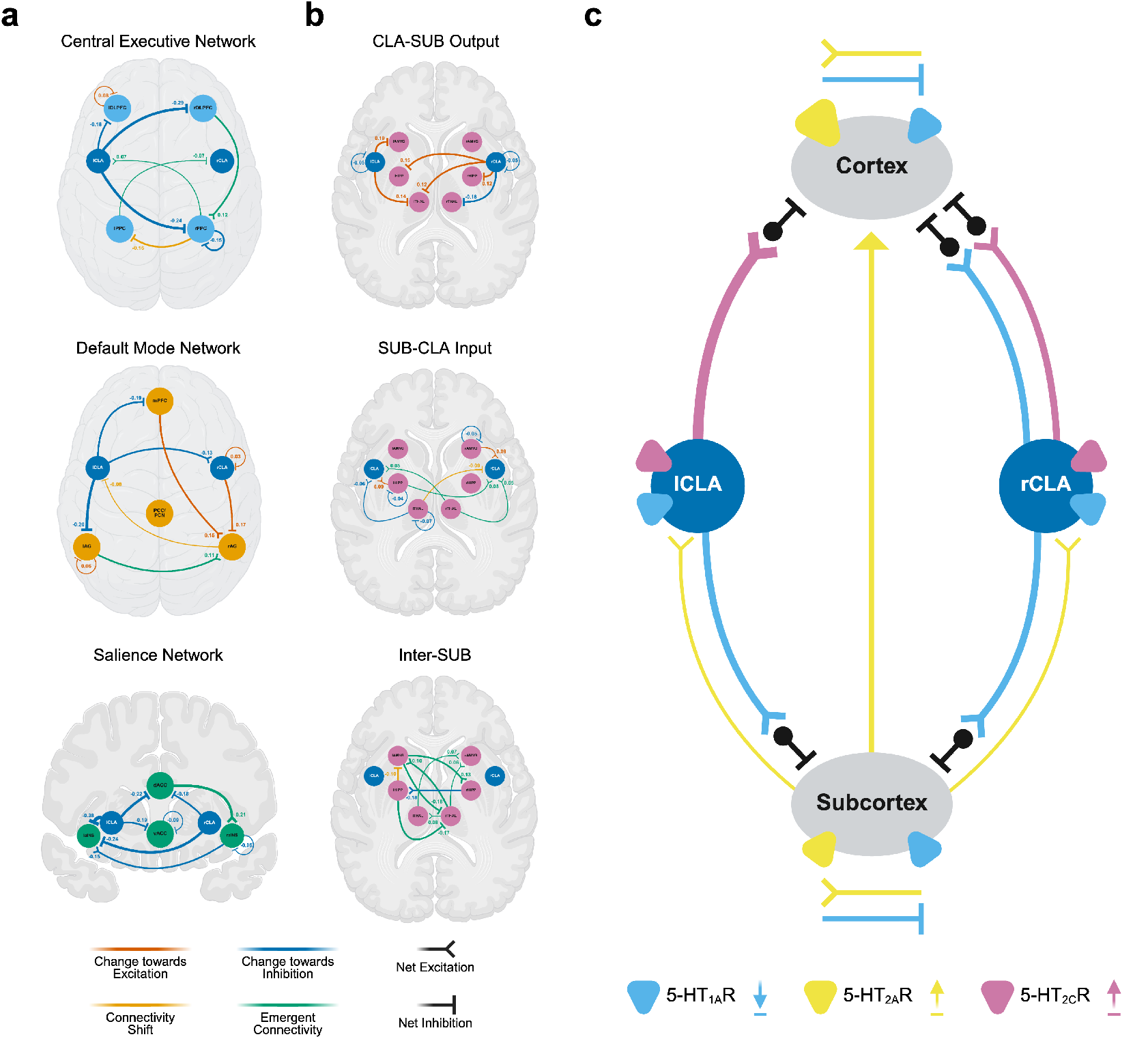
Changes in effective connectivity under psilocybin. **a** Triple-network connectivity changes involving the claustrum and within-network alterations. The magnitude of EC change is reflected by the arrow thickness in incremenents of 0.20 Hz.**b** Subcortical connectivity changes, including claustral outputs, subcortical inputs, and inter-subcortical influences. Here the magnitude of EC change is reflected by the arrow thickness in incremenents of 0.10 Hz. **c** Schematic of 5-HT receptor–mediated shifts in excitatory–inhibitory signalling between CLA hemispheres, cortex, and subcortex. 5-HT_1A_Rs are generally inhibitory, while 5-HT_2_ subtypes are excitatory. Based on our EC findings and the existing literature we propose that 5-HT receptor subtypes may mediate differential aspects of the EC shifts observed here, with 5-HT_2C_Rs upregulating claustral inhibibition of the cortex, which was more pronounced from the left CLA, while 5-HT_1A_Rs mediate claustro-subcortical, and node-specific decreases in inhibition. 5-HT_2A_Rs increase subcortical excitation of both claustra, and similarly increase information flow to the cortex, which was not characterized here. Lastly, 5-HT_1A_ and 5-HT_2A_Rs differentially shift dynamics within the cortex and subcortex. Receptor icon size reflects relative expression levels in the respective area.

Taken together, psilocybin selectively modulated claustro-cortical signaling, showing a strong left-lateralized upregulation of claustral inhibition, especially pronounced for the CEN and DMN. Right CLA modulation was more nuanced, showing no effects on CEN nodes, decreased inhibition of the right AG, and SN node-specific upregulation of inhibition.

### Psilocybin modulates cortico-claustral input and within-network signaling

Changes in cortical input to the CLA under psilocybin were less pronounced than the upregulation of claustro-cortical inhibition (see Fig. 4**a**). In the CEN, emergent excitation from the bilateral PPC to contralateral claustra became apparent, while in the DMN, only an excitatory-to-inhibitory switch from the right AG to left CLA was observed, all at comparable strengths. In the SN, no changes in CLA input were detected.

Psilocybin more strongly shifted within-network EC. In the CEN, an excitatory influence from the right DLPFC to the right PPC emerged, while the right PPC switched from excitation of the left PPC to inhibition. Additionally, a slight decrease and increase in self-inhibition were estimated for the left DLPFC and right PPC, respectively. In the DMN, excitation to the right AG increased, driven by stronger mPFC input and emergent input from the left AG, which displayed a decrease in self-inhibition. In the SN, excitation from the right aINS to the left aINS decreased, while a strong excitatory influence from the dACC to the right aINS emerged.

In summary, psilocybin modulated cortico-claustral input selectively from parietal nodes of the CEN and DMN. Within-network changes were characterized by enhanced or emergent frontal excitation to, and shifts between, right parietal and insular nodes.

### Psilocybin reduces claustro-subcortical inhibition while enhancing subcortical input

In contrast to its effects on claustro-cortical signaling, psilocybin decreased CLA inhibition of subcortical regions, with more connections modulated from the right CLA (see Fig. 4**b**). The left CLA disinhibited the left AMYG and left THAL, while the right CLA disinhibited the bilateral HIPP and left THAL but increased inhibition of the right THAL, all at comparable strengths. Additionally, both claustra displayed an equal slight increase in self-inhibition.

This subcortical disinhibition was accompanied by marked increases in subcortical input to the CLA. Both claustra received emergent excitation from the right THAL. Tn contrast the left THAL decreased its excitatory influence, reducing inhibition of the left CLA and switching from excitation to inhibition of the right CLA. Both claustra also received increased excitation from the left HIPP, which was emergent to the right CLA. The right AMYG displayed increased excitation only of the right CLA. No change in left AMYG signaling was detected. All shifts in subcortical input to the CLA were of comparable magnitude, including slightly increased self-inhibition estimated for the right AMYG, left HIPP, and left THAL.

Overall, psilocybin shifted CLA–subcortex dynamics toward decreased claustral inhibition of subcortical regions and enhanced excitatory input to both claustra, with a divergent pattern observed for thalamic hemispheres, which strongly contrasts the shifts observed for cortical networks.

### Psilocybin strongly modulates subcortical signalling

Psilocybin profoundly altered subcortex dynamics, marked by various emergent influences between subcortical regions (see Fig. 4**b**). Markedly, three excitatory influences from the right THAL emerged, targeting the bilateral AMYG and left THAL, while the left THAL started exciting the right AMYG. From the left AMYG, an excitatory influence on the right HIPP, and inhibition of the right THAL emerged. The right HIPP decreased its excitation of the left HIPP, while the left HIPP switched from excitation to inhibition of the left AMYG and displayed emergent inhibition of the right THAL. No change in right AMYG signalling was estimated. Notably, changes in EC between subcortical regions displayed differences in magnitude, which were especially pronounced for hippocampal and right amygdala signalling.

Taken together, psilocybin strongly enhanced subcortical communication between all three regions, with most signals emerging from the THAL, but stronger modulation of hippocampal and left lateralized AMYG signaling.

### Changes in claustro-cortical and subcortical signaling are associated with psychedelic subjective effects

We next examined whether alterations in claustro-cortical and claustro-subcortical signaling were associated with differential aspects of psychedelic subjective effects, using PEB-based analyses with the MEQ30 and 11D-ASC composite scores as covariates (see Table 1)

**Table 1:** Effective connectivity changes associated with self-report measures of psychedelic subjective effects.

| Scale | Network | Connection | EC Change | Net EC (Hz) | $\beta$ Coefficient |
| --- | --- | --- | --- | --- | --- |
| MEQ30: Total | CEN | CLA_L → DLPFC_L | ↑ Inhibition | -0.56 | 0.239 |
|  |  | CLA_L → DLPFC_R | ↑ Inhibition | -0.68 | 0.184 |
|  |  | PPC_R → PPC_L | Excitation → Inhibition | -0.05 | -0.120 |
|  | SN | CLA_R → aINS_L | ↑ Inhibition | -0.97 | 0.147 |
|  | SUB | CLA_L → AMYG_L | ↓ Inhibition | -0.12 | -0.142 |
|  |  | HIPP_L → THAL_R | Emergent Inhibition | -0.17 | 0.098 |
|  |  | THAL_R → AMYG_R | Emergent Excitation | 0.10 | 0.073 |
|  |  | AMYG_R | ↑ Self-Inhibition | -0.55 | 0.120 |
|  |  | HIPP_L | ↑ Self-Inhibition | -0.72 | -0.148 |
| 11D-ASC: Associative | CEN | CLA_L → DLPFC_L | ↑ Inhibition | -0.56 | 0.228 |
|  |  | CLA_L → DLPFC_R | ↑ Inhibition | -0.68 | 0.301 |
|  |  | CLA_L → PPC_R | ↑ Inhibition | -0.73 | 0.245 |
|  |  | PPC_R → PPC_L | Excitation → Inhibition | -0.05 | -0.089 |
|  | DMN | AG_R → CLA_L | Excitation → Inhibition | -0.03 | 0.051 |
|  | SUB | CLA_L → AMYG_L | ↓ Inhibition | -0.12 | -0.145 |
|  |  | HIPP_R → HIPP_L | ↓ Excitation | 0.15 | -0.126 |
|  |  | HIPP_L | ↑ Self-Inhibition | -0.72 | -0.174 |
|  |  | THAL_L | ↑ Self-Inhibition | -0.38 | -0.125 |
| 11D-ASC: Sensory | CEN | CLA_L → DLPFC_R | ↑ Inhibition | -0.68 | 0.142 |
|  | DMN | AG_L | ↓ Self-Inhibition | -0.30 | -0.301 |
|  | SUB | HIPP_L → AMYG_L | Excitation → Inhibition | -0.03 | 0.112 |
|  |  | AMYG_R | ↑ Self-Inhibition | -0.55 | 0.121 |
|  |  | HIPP_L | ↑ Self-Inhibition | -0.72 | -0.123 |
| 11D-ASC: Negative | CEN | PPC_R | ↑ Self-Inhibition | -0.47 | 0.222 |
|  | DMN | CLA_L → mPFC | ↑ Inhibition | -0.92 | -0.168 |
|  | SUB | CLA_R → THAL_R | ↑ Inhibition | -0.54 | -0.139 |
|  |  | HIPP_L → CLA_L | ↑ Excitation | -0.14 | 0.051 |
|  |  | THAL_R → THAL_L | Emergent Excitation | 0.08 | 0.091 |
|  |  | THAL_L | ↑ Self-Inhibition | -0.38 | -0.119 |

The total mystical experience (MEQ30: Total) showed associations with CEN, SN, and subcortical connections. Increased inhibition from the left CLA to the bilateral DLPFC showed the strongest positive association, followed by increased inhibition from the right CLA to the left aINS, increased right AMYG self-inhibition, and emergent inhibition from the left HIPP to the right THAL. Negative associations were observed with increased left HIPP self-inhibition, reduced inhibition from the left CLA to the left AMYG, and an excitatory-to-inhibitory switch from the right to left PP

Similarly, 11D-ASC: Associative scores implicated the CEN, DMN, and subcortex, and were most strongly positively associated with increased inhibition from the left CLA to bilateral DLPFC and right PPC, followed by an excitatory-to-inhibitory switch from the right AG to the left CLA. Negative associations were strongest with the subcortex, including increased HIPP self-inhibition, reduced inhibition from the left CLA to the left AMYG, decreased right-to-left HIPP excitation, stronger left THAL self-inhibition, and lastly, a right-to-left PPC excitatory-to-inhibitory switch.

11D-ASC: Sensory scores again showed associations with CEN, DMN, and subcortex, with the strongest positive association found for increased left CLA to right DLPFC inhibition, an excitatory-to-inhibitory switch from the left HIPP to left AMYG, and increased right AMYG self-inhibition. A strong negative association was found with decreased left AG self-inhibition, followed by increased left HIPP self-inhibition.

Finally, 11D-ASC: Negative scores implicated the same networks. Increased right PPC self-inhibition showed the strongest positive association, followed by emergent right-to-left THAL excitation, and increased excitation from the left HIPP to left CLA. 11D-ASC: Negative scores showed the strongest negative association with increased left CLA to mPFC inhibition, followed by increased inhibition from the right CLA to right THAL and increased left THAL self-inhibition.

In summary, we identified various connectivity changes that were associated with differential aspects of psychedelic subjective effects, positively associating increased inhibition from the CLA to frontal cortical network nodes to all measures, while increased claustro-subcortical inhibition only displayed negative associations. Furthermore, within-network and subcortical signaling changes showed a more nuanced pattern of pathway-specific associations, predominantly linking subcortical signaling modulation to overall mystical, sensory, and negative effects, with uniformly negative associations observed with sensory effects.

### Evaluation of small region confound correction

We employed small region confound correction (SRCC) to control for partial volume effects of surrounding regions on the extracted CLA BOLD time-series. To do so, we created so-called flanking region masks based on the INS and PUT and used them as regressors in a linear regression model of the CLA BOLD time series (see Fig. 5a). In this step, we calculated the mean coefficient of determination (R^2^) to determine the variance explained in the CLA time-series by flanking regions. The variance explained was much greater in HCP7T (Left Flank: mean R^2^ = 0.473, SD = 0.270; Right Flank: mean R^2^ = 0.453, SD = 0.273) than for HCP3T (Left Flank: mean R^2^ = 0.080, SD = 0.095; Right Flank: Left Flank: mean R^2^ = 0.067, SD = 0.077) and PC3T (Left Flank: mean R^2^ = 0.074, SD = 0.223; Right Flank: Left Flank: mean R^2^ = 0.020, SD = 0.088).

**Fig. 5:**
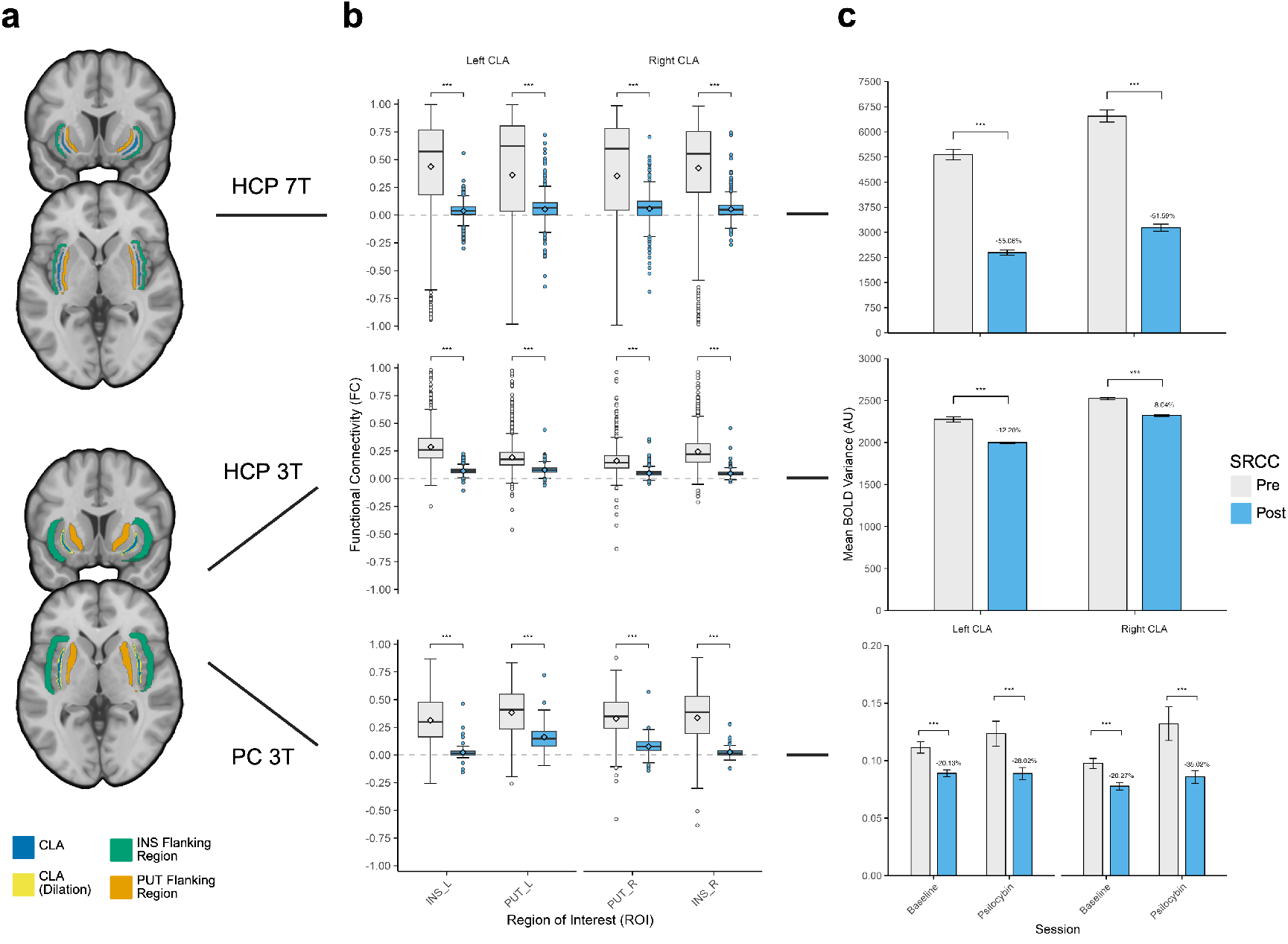
Small Region Confound Correction (SRCC) Metrics. **a** SRCC masks are adjusted to functional voxel size, corresponding to two functional voxels (3mm at 7T; 6mm at 3T) in size and distance to the CLA. At 3T, CLA masks were dilated by 0.75mm to ensure reliable extraction at the larger voxel size. **b** SRCC reliably reduced FC of the CLA to the surrounding Putamen (PUT) and Insula (INS) towards zero across datasets, confirming significant removal of potential partial volume effects. **C** By definition, SRCC will reduce CLA blood oxygen level-dependent (BOLD) variance by removing any variance explained by surrounding regions. While the reduction of FC to surrounding was comparable across datasets, the magnitute of CLA BOLD variance reduction varied substantially between datasets. Additionally, no significant difference in CLA BOLD variance was observed between the baseline and psilocybin scans in PC3T. Box plots span the Interquartile Range (IQR), with median line, mean diamond, whiskers within bounds ±1.5 IQR, and outlier dots. Bar charts display mean ± SEM. Significance testing is based on the Mann-Whitney U-test. P-values are adjusted via Bonferroni Correction (** = adj. p *<* 0.01, *** = adj. p *<* 0.001). AU - arbitrary unit

To evaluate the success of this correction, we calculated the functional connectivity (FC) between the CLA and the ipsilateral PUT and insula INS pre- and post-SRCC (see Fig.5b). In HCP7T, the mean FC between each CLA to the ipsilateral INS and PUT was significantly reduced towards zero (Left INS: Mean Difference (MD) = -0.399, SD = 0.413, Cohen’s dz = -0.967, adj. p *<* 0.001; Left PUT: MD = -0.309, SD = 0.515, dz = -0.600, adj. p *<* 0.001; Right INS: MD = -0.371, SD = 0.420, dz = -0.882, adj. p *<* 0.001; Right PUT: MD = -0.295, SD = 0.491, dz = -0.602, adj. p *<* 0.001). The same goes for HCP3T (Left INS: MD = -0.216, SD = 0.135, dz = -1.599, adj. p *<* 0.001; Left PUT: MD = -0.114, SD = 0.096, dz = -1.183, adj. p *<* 0.001; Right INS: MD = -0.197, SD = 0.124, dz = -1.584, adj. p *<* 0.001; Right PUT: MD = -0.112, SD = 0.088, dz = -1.280, adj. p *<* 0.001), and PC3T (Left INS: MD = -0.290, SD = 0.231, dz = -1.259, adj. p *<* 0.001; Left PUT: MD = -0.221, SD = 0.182, dz = -1.216, adj. p *<* 0.001; Right INS: MD = -0.311, SD = 0.269, dz = -1.156, adj. p *<* 0.001; Right PUT: MD = -0.253, SD = 0.193, dz = -1.310, adj. p *<* 0.001).

Lastly, to evaluate the magnitude of SRCC on the extracted CLA BOLD time-series, we compared CLA BOLD variance pre- and post-SRCC (see Fig. 5c). In HCP7T, SRCC reduced CLA BOLD variance by over 50% (Left CLA: MD = -2922.638, SD = 3526.500, dz = - 0.829, adj. p *<* 0.001; Right CLA: MD = -3331.552, SD = 4140.340, dz = -0.805, adj. p *<* 0.001). In HCP3T, SRCC reduced CLA BOLD variance by up to 12% (Left CLA: MD = -277.627, SD = 1868.628, dz = -0.829, adj. p *<* 0.001; Right CLA: MD = -202.942, SD = 653.027, dz = -0.311, adj. p *<* 0.001). In PC3T, SRCC reduced CLA BOLD variance by up to 28% for the baseline scan (Left CLA: MD = -0.022, SD = 0.029, dz = -0.784, adj. p *<* 0.001; Right CLA: MD = -0.020, SD = 0.021, dz = -0.931, adj. p *<* 0.001), and up to 35% for the psilocybin scan (Left CLA: MD = - 0.035, SD = 0.059, dz = -0.528, adj. p *<* 0.001; Right CLA: MD = -0.046, SD = 0.087, dz = -0.529, adj. p *<* 0.001). No significant difference in CLA BOLD variance was detected pre-SRCC (Left CLA: MD = 0.012, SD = 0.080, dz = 0.150, adj. p = 1.000; Right CLA: MD = 0.035, SD = 0.107, dz = 0.323, adj. p = 0.231), and post-SRCC (Left CLA: MD = 0.000, SD = 0.038, dz = -0.004, adj. p *<* 0.469; Right CLA: MD = -0.008, SD = 0.045, dz = 0.180, adj. p = 1.000).

Overall, SRCC reduced FC between the CLA and surrounding structures, and therefore potential partial volume effects, similarly effective in all datasets, across resolutions. However, the loss in CLA BOLD variance was markedly greater at 7T, probably due to greater proximity between flanking region and CLA voxels, contrary to a higher resolution less susceptible to partial volume effects.

## Discussion

Here, we present the first human in vivo characterization of EC between the CLA, major cortical networks, and subcortical regions. Employing spDCM on rs-fMRI data from two independent datasets, at three different voxel sizes, enabled us to extend the current CLA literature by estimating a select input-output organization of the human CLA at the regional level, providing a quantification of net excitatory and inhibitory influences, as well as modulation of these signalling pathways under psilocybin.

Methodologically, our findings support sound imaging of the CLA in humans. The small anatomical size of the CLA remains below the currently possible resolutions, even at high MRI field strengths (Kapakin, 2011), rendering signal contamination a very probable risk, for which SRCC provides a simple yet very conservative approach. We show that SRCC reduces FC of the CLA to surrounding areas in a similar manner across spatial resolutions. However, the loss of CLA BOLD variance was substantial at 7T, although less susceptible to partial volume effects. Importantly, the directionality of signal correlation (e.g., FC) between the CLA, PUT, and INS remains to be determined. Thus, a reduction of FC may not only reflect partial volume effects, but also functionally relevant signalling from the CLA to surrounding structures, vice versa, or both. Here, application of SRCC did not substantially affect DCM outcomes, which, most strikingly, still provided EC estimates for the aINS, exemplifying the non-direct mapping of FC to EC (Novelli et al., 2024). However, in CLA studies employing FC as a measure that include the SN or striatum, employment of SRCC holds inevitable implications for FC outcomes. Accurate CLA segmentation remains a challenge, with manual approaches presenting a time-consuming endeavour, especially at large sample sizes (Kang et al., 2025). Data sharing efforts of pre-made CLA masks are of great value in making human CLA imaging more feasible and accessible (Coates and Zaretskaya, 2024; Rodríguez-Vidal et al., 2024; Stewart et al., 2024). However, future research would greatly benefit by employing subject-specific CLA masks to increase extraction accuracy, for which automatic deep-learning approaches have been developed (Berman et al., 2020; Li et al., 2021; Mauri et al., 2025).

On a higher level, our findings translate structural animal data, replicating widespread bidirectional connectivity of the CLA to cortical network nodes (Jackson et al., 2020), and confirm recent transcriptomics findings in primates, extending the previously suggested unidirectional connectivity of subcortical regions to bidirectional influences between the CLA, amygdala, hippocampus, and thalamus (Lei et al., 2025). Furthermore, we confirm and extend diffusion-weighted imaging and FC data in humans (Milardi et al., 2015; Rodríguez-Vidal et al., 2024; Wendt et al., 2024), providing a directed assessment of CLA signalling relevant cortical network dynamics (Torgerson et al., 2015), and excitatory inter-claustral coupling structurally mediated via the corpus callosum (Arrigo et al., 2017). Overall, we show that claustral output induces strong and uniform inhibition of all cortical network nodes, and to a weaker extent of subcortical targets, all while integrating predominantly excitatory input from these same areas.

The inhibitory influence of the CLA aligns with cellular and electrophysiological animal data, suggesting predominant mapping of claustral efferents onto cortical inhibitory interneurons (de la Torre-Martínez et al., 2025; McBride et al., 2023). Albeit weaker than cortical inhibition, to our knowledge, no direct investigation of claustral signalling to subcortical regions reports an analogous cellular mapping. Our findings suggest that the CLA may induce subcortical inhibition via a similar FFI mechanism as identified in the cortex. These findings hold important implications for current functional theories of CLA signalling. In the cortex, claustro-cortical inhibition has been proposed to underlie cross-frequency-coupling (CFC) linking claustral low-frequency oscillations to higher cortical rhythms to synchronize functionally-connected but distal cortical regions (Vidyasagar and Levichkina, 2019). Intriguingly, our EC estimates closely match the ACC-CLA input (1-10 Hz) and CLA-ACC output (50-80 Hz) frequencies directly measured in mice (**?**). The more recent NICC model postulates a similar functional role of claustro-cortical inhibition, suggesting that frontal regions, such as the mPFC and ACC, direct the CLA to instantiate cortical network signatures in a task-dependent manner (Madden et al., 2022). While we did not find evidence for stronger input from frontal nodes, we did find a weaker inhibitory influence of the CLA on the ventral subdivision of the ACC than on the dACC, with the vACC generally not considered a key node of the SN (Uddin et al., 2019). Inhibitory control of the CLA over the SN was a more unexpected finding, and particularly the strong inhibition of the right CLA over the right aINS is intriguing, given that the right aINS is considered the key output hub of the SN involved in switching between the CEN and DMN (Menon and Uddin, 2010; Uddin, 2015). We found that claustral inhibition of the DMN was somewhat stronger than for the CEN and SN at rest, potentially reflecting sustained maintenance of a DMN state. Interestingly, the PCC/PCN region was found to be the only region inducing inhibition of the CLA at rest, suggesting a differential influence of cingulate signaling to the CLA from anterior and posterior nodes, as part of the SN and DMN, respectively.

The latest functional proposal suggests that the CLA not only coordinates synchronization of cortical networks but is involved in synchronized brain states as a whole, by inducing cortical downstates characterized by neural suppression and low-frequency activity (Do et al., 2024). Again, cortico-claustral inhibition may be integral to such function, but the authors also highlight the differential contributions of thalamic and hippocampal oscillations in regulating the full spectrum of cortical synchrony (Gent et al., 2018; Maingret et al., 2016; Peyrache et al., 2011; Poulet et al., 2012). In that regard, our findings may additionally hold functional implications on how the CLA could be involved in brain synchrony by also coordinating subcortical rhythms. The implications of CLA signalling to the amygdala are more elusive. However, recent animal data implicate amygdala-CLA signalling in social stress-related domains (Froesel et al., 2024; Niu et al., 2022; Tanuma et al., 2022).

It is important to consider that our findings can only be interpreted in the context of resting-state acquisition. Our EC estimates oppose recent task-based findings suggesting excitatory signalling to key nodes of the same networks (Stewart et al., 2025). For the same reason, we probably did not identify more pronounced frontal signalling to the CLA, which may require task-engagement (Madden et al., 2022). Still, the cellular mapping of the CLA to cortical regions is highly complex, depending on cortical layer and targeted cell type (de la Torre-Martínez et al., 2025; McBride et al., 2023; Wang et al., 2023), entertaining the possibility that the CLA also may induce excitatory EC in a context-dependent manner. The differential functional relevance of CLA excitatory and inhibitory output effects is something future research should address.

We also identified pronounced modulation of CLA EC under psilocybin, specifically marked by a lateralized upregulation of claustro-cortical inhibition of the DMN and CEN, which was more balanced for the SN. Contrary to these cortical effects, we observed a disinhibition of subcortical regions and increased input signalling to the CLA and between subcortical regions. Association analysis with subjective effect scores predominantly implicated increased claustral inhibition of frontal cortical nodes, especially the DLPFC, in mystical and associative scores, while subcortical signalling changes displayed a more nuanced pattern, displaying positive associations with mystical, negative associations with associative, and mixed associations with sensory and negative effects.

Divergent modulation of claustral inhibition of cortical and subcortical targets may be attributable to differential expression and, therefore, functional localization of 5-HT receptor subtypes. As such, rodent evidence suggests that psychedelics upregulate glutamate receptor–mediated EPSPs on claustral projection neurons in a 5-HT_2C_R-dependent manner, whereas 5-HT_1A_R activation inhibits CLA output (Anderson et al., 2024). This way, claustro-cortical FFI may be enhanced via HT_2C_R activation, whereas claustro-subcortical projections are inhibited due to differential expression of HT_1A_R. Investigations into the target-specific localization of 5-HT receptor subtypes on claustral efferents would greatly benefit addressing this possibility. Similarly, the mechanisms explaining the observed lateralization of claustro-cortical inhibition remain elusive. Notably, lateralized modulation of CLA FC to the DMN and fronto-parietal network under psilocybin has previously been shown in humans, which was differentially associated with modularity and integrity of these networks (Barrett et al., 2020). In primates, psilocybin has been shown to increase FC between the CLA, ACC, PFC, and AG, which, however, did not distinguish CLA hemispheres (Bagdasarian et al., 2024). While it is not straightforward to directly relate these findings to ours, due to the many-to-one mapping of FC to EC (Novelli et al., 2024), in light of our evidence, these findings converge on lateralized modulation of CLA connectivity to exactly these network nodes. A hemispheric asymmetry in claustro-cortical inhibition may manifest in decreased within-network FC characteristic of psychedelic action by differentially modulating select nodes, and thus reducing network synchrony (Siegel et al., 2024; Yu et al., 2024). Still, contrary to the current conceptualisation of the CCC model, which posits a two-hit mechanism of 5-HT_2A_R-dependent modulation of frontal cortico-claustral input and claustro-cortical output (Doss et al., 2022), we did not find changes in frontal cortical input to the CLA. Taken together, we propose a different framework of psychedelic modulation, integrating the hemispheric and hierarchical asymmetry in claustral signaling modulation with 5-HT receptor mechanisms, considering our findings and the literature reviewed here (see Fig. 4**c**).

The clinical implications of our findings are manifold. Bridging the functional role of the CLA in cortical network coordination (Madden et al., 2022), and a disruption of this mechanism under psychedelic action (Doss et al., 2022), the triple network model provides a transdiagnostic framework, positing that differentially aberrant signalling between the CEN, DMN, and SN may manifest in distinct pathological phenotypes of dysfunctional cognitive control, self-referential, and salience processing (Menon, 2019, 2011). Altered dynamics between these networks have been reported across disorders, including affective disorders (Wang et al., 2020; Zheng et al., 2015), Alzheimer’s disease (Li et al., 2019; Meng et al., 2022), schizophrenia (Menon et al., 2023; Supekar et al., 2019), obsessive-compulsive disorder (Fan et al., 2017; Li et al., 2025), and chronic pain (De Ridder et al., 2022), coinciding with disorders increasingly implicating the CLA (Ayyildiz et al., 2023; Schinz et al., 2024; Zhang and Zamponi, 2024). Psychedelics have been repeatedly shown to modulate connectivity between these networks (Melani et al., 2025; Zhang et al., 2025), coinciding with therapeutic efficacy across multiple indications (Andersen et al., 2021; Heifets and Olson, 2024; Ko et al., 2022). Thus, CLA may represent a promising subcortical target mediating aberrant network signalling and the therapeutic efficacy of psychedelics (Benarroch, 2021; Nichols et al., 2017).

The current study is still subject to some limitations. Here, we did not adjust SRCC to the HCP3T dataset, which had a functional voxel size of 2 mm, rendering this data more susceptible to partial volume effects. Furthermore, we did not directly inspect the overlap of our CLA mask with T1w images across all participants due to the sheer number of subjects included. It is important to note that the HCP 3T and 7T datasets are not independent. Furthermore, the HCP3T data includes over 150 twins and provides a high spatial resolution. Thus, outcome convergence between the HCP7T and 3T data should be interpreted with caution and may be attributable to a shared subject pool and close spatial resolution. Here, we estimated four separate models of CLA EC to cortical networks and the subcortex. A large-scale DCM integrating all ROIs would require greater computational power and time, especially given the large sample size used in this study, all while increasing model complexity and the number of parameters to be estimated. Many of the additional EC estimates would not have been of interest to our CLA-centered hypothesis. For the same reason, we did not discuss within-network EC estimates at rest. We did not directly test the functional implications of our findings, which remain to be explicitly empirically investigated. Similarly, we did not investigate EC between the CLA, sensory, and motor cortices, all of which show strong functional and structural links to the CLA. It remains to be investigated if similar EC dynamics exist with sensory and motor networks.

To conclude, we present a thorough characterization of the effective connectivity of the claustrum to the central executive, default mode, and salience network, as well as subcortical regions such as the amygdala, hippocampus, and thalamus in humans at rest. Our findings suggest that the claustrum exerts strong inhibitory control over cortical network nodes, which may subserve network synchrony. Similarly, this inhibitory control extends to the subcortex, albeit at a weaker strength, opening new theoretical avenues of claustral functional influence on subcortical dynamics. Similarly, the claustrum integrates a wide array of inputs from all regions investigated, strengthening its role as an integrative relay-hub. Furthermore, we showed that psilocybin asymmetrically modulates claustral signaling, displaying hemispheric lateralization in upregulation of cortical inhibition, and a hierarchical asymmetry by in turn reducing inhibition of the subcortex and increasing signaling to the claustrum and between these regions. Overall, many questions remain to be addressed by future research, including the optimization of claustral imaging in humans, receptor-informed characterization of claustral signaling modulation, and the therapeutic implications of claustro-cortical and subcortical dynamics.

## Methods

### Datasets and subjects

#### Human Connectome Project

The data analyzed in this study were derived from two different datasets. The Human Connectome Project (HCP) dataset is part of the WU-Minn HCP 1200 Subjects Data Release provided by the Human Connectome Project, WU-Minn Consortium (Principal Investigators: David Van Essen and Kamil Ugurbil; 1U54MH091657) funded by the 16 NIH Institutes and Centers that support the NIH Blueprint for Neuroscience Research; and by the McDonnell Center for Systems Neuroscience at Washington University. The HCP dataset officially comprises behavioral and demographic data from 1206 healthy young adults (Age: 22–35 years), spanning 3T and 7T fMRI, magnetoencephalography (MEG), and genotyping data. Exclusion criteria included a history of psychiatric, neurological, or neurodevelopmental disorders or substance abuse (see (Van Essen et al., 2013) for details). 3T fMRI data is available for 1113 subjects (Gender: 507 males, 606 females) (Luo et al., 2019), of which a subset of 184 subjects additionally underwent 7T fMRI. Here, we analyzed the entire resting-state (rs-fMRI) data of the HCP dataset available to us at 7T (N = 184) and 3T (N = 1096). All but some subjects underwent up to four rs-fMRI scanning sessions at both field strengths, resulting in a total of 722 and 4223 scans analyzed at 7T and 3T, respectively. For further information on the study protocol and subjects, please refer to the WU-Minn HCP 1200 Subjects Data Release Reference Manual and see (Elam et al., 2021).

#### PsiConnect

The PsiConnect (PC3T) dataset comprises neuroimaging and psychometric data from 62 healthy and psychedelicnaïve adults (Age: 18 – 55, Gender: 30 females, 31 males, 1 non-binary). Exclusion criteria included a history of psychiatric disorders or suicidality, a 5-year history of substance and/or alcohol use disorder, first-degree relatives with a diagnosed psychotic disorder, a history of major neurological disorders including stroke and epilepsy, use of hallucinogens within the last six months, a formal meditation practice in the last 6 months or extensive previous exposure to mindfulness meditation, and contraindicated medications. The study protocol was approved by the Monash University Human Research Ethics Committee and registered with the Australian New Zealand Clinical Trials Registry under the registration number ACTRN12621001375842.

Subjects were randomly assigned to one of two matched groups, with half of the participants undergoing an 8-week mindfulness-based cognitive therapy (MBCT) program pre scanning. All subjects underwent a multimodal naturalistic imaging protocol employing fMRI and subsequent electroencephalography (EEG) over two sessions, at baseline and under the acute influence of 19 mg of oral synthetic psilocybin, two weeks apart. The imaging protocol consisted of four blocks across fMRI and EEG data acquisitions, employing eyes closed resting-state, guided meditation and music listening, and eyes-opened movie watching. Scores on multiple psychometric self-rating questionnaires were collected at various time-points across, between, and after imaging sessions. For further information on the PsiConnect study protocol see (Novelli et al., 2025) and (Stoliker et al., 2025). Here, we analyzed only the resting-state block of the fMRI data at baseline and under psilocybin, including a total of 59 subjects. Three subjects were excluded from the analysis due to significant outlier image quality metrics identified via MRIQC. We pooled all subjects together, not distinguishing between the meditation and non-meditation groups, due to negligible effects of meditation training on fMRI measures (Stoliker et al., 2025).

We included data from two psychometric self-report measures of psychedelic subjective effects, namely the total scores of the 30-item revised Mystical Experience Questionnaire (MEQ30) (Barrett et al., 2015) and composite scores of the 11 Dimensions of Altered States of Consciousness (11D-ASC) scale (Studerus et al., 2010). The MEQ30: Total score was calculated per subject by summing the raw scores of all four factors and dividing them by 150 (maximum possible MEQ30 score), to obtain the total score expressed as the percentage of the maximum score possible (Barrett and Griffiths, 2018). The MEQ30: Total score has been shown to have the greatest predictive validity for capturing the underlying latent mystical experience and is preferable when not testing factor-specific hypotheses (Barrett et al., 2015).

For the 11D-ASC scale, a total score per subject was calculated as the percentage of the maximum score possible for each of the 11 factors. Three composite scores were calculated by summing and averaging relevant factor scores per subject. The first composite score was termed 11D-ASC: Associative, based on the first three factors: (1) Experience of Unity, (2) Spiritual Experience, and (3) Blissful State. The second composited score was termed 11D-ASC: Sensory, based on the three factors: (8) Complex Imagery, (9) Elementary Imagery, and (10) Audio-Visual Synaesthesia. The third composite score was termed 11D-ASC: Negative, based on the two factors: (6) Impaired Control and Cognition, and (7) Anxiety. Rationale for computing these composite scores stems from significantly high correlations between the factors of each composite score observed in the PC3T sample (Stoliker et al., 2025). Additionally, factors of each composite score load exclusively onto one of the five higher-order dimensions of the ASC scale (5D-ASC), with the 11D-ASC: Associative mapping onto Oceanic Boundlessness, 11D-ASC: Sensory onto Visionary Destructuralization, and the 11D-ASC: Negative onto Dread of Ego Dissolution (Hirschfeld and Schmidt, 2021). Data was missing for two subjects for the MEQ30: Total, three subjects for the 11D-ASC: Associative, five subjects for the 11D-ASC: Sensory, and eight subjects for the 11D-ASC: Negative scores, which were thus excluded from the behavioral association analysis.

#### Data acquisition and preprocessing

HCP7T rs-fMRI data were acquired on a Siemens Magnetom 7T MR scanner at the Center for Magnetic Resonance (CMRR) at the University of Minnesota in Minneapolis, MN, using a Nova32 32-channel Siemens receive head coil with an incorporated head-only transmit coil that surrounds the receive head coil from Nova Medical (Sequence: Gradient-echo EPI; TR = 720 ms; TE = 33.1 ms; Flip Angle = 52°; FOV = 208 x 180 mm (RO x PE); Matrix = 104 x 90 (RO x PE); Functional Voxel Size = 1.6 mm isotropic; Multiband factor = 8; Echo Spacing = 0.58 ms; BW = 2290 Hz/Px). Oblique axial acquisitions alternated between phase encoding in a posterior-to-anterior (PA) direction in runs 1 and 3, and an anterior-to-posterior (AP) phase encoding direction in runs 2 and 4.

HCP 3T rs-fMRI data were acquired on a on a customized Siemens 3T “Connectome Skyra” housed at Washington University in St. Louis, using a standard 32-channel Siemens receive head coil and a “body” transmission coil (Sequence: Gradient-echo EPI; TR = 1000 ms; TE = 22.2 ms; Flip Angle = 45°; FOV = 208 x 208 mm (RO x PE); Matrix = 130 x 130 (RO x PE); Functional Voxel Size = 2.0 mm isotropic; Multiband factor = 8; Echo Spacing = 0.58 ms; BW = 1924 Hz/Px). Oblique axial acquisitions alternated between phase encoding in a right-to-left (RL) direction in runs 1 and 3, and a left-to-right (LR) phase encoding direction in runs 2 and 4.

During HCP MRI acquisition, subjects kept their eyes open, instructed to maintain a relaxed fixation on a projected bright crosshair on a dark background. For most but not all subjects, acquisition duration lasted for approximately 14:30 min, acquiring 1200 frames, and 16 minutes, acquiring 900 frames, per run at 7T and 3T, respectively. The first 5 frames were omitted from analysis.

PC3T rs-fMRI data were acquired at the Monash Biomedical Imaging facility, Monash University, Australia using a Siemens Magnetom Skyra 3T MRI scanner equipped with a 32-channel receive head coil (Sequence: Multi-echo, multi-band accelerated EPI (mbep2d); TR = 910 ms, TE1 = 12.60 ms, TE2 = 29.23 ms, TE3 = 45.86 ms, TE4 = 62.49 ms; FOV = 206 mm; Functional Voxel Size = 3.2 mm isotropic, Multiband factor = 4; Image Acceleration factor (PE) = 2 and 3). Functional data was acquired in an RL phase encoding direction. Participants kept their eyes closed during acquisition. Resting-state blocks lasted for approximately 8 minutes, acquiring 505 frames per session, of which the first 5 were omitted from analysis.

All datasets were provided in preprocessed form, for which different pipelines were employed. HCP dataset was preprocessed using a custom minimal preprocessing pipeline (see (Glasser et al., 2013)) for details). PC3T data was preprocessed using the fMRIprep 22.0.2 pipeline (Esteban et al., 2019), and image quality control was conducted using MRIQC BIDS (Esteban et al., 2017) (see (Novelli et al., 2025) for details).

#### Regions of interest

High-resolution masks for the left and right CLA, insula, and putamen were made available by (Stewart et al., 2024). These mean masks were manually created based on 7T-acquired T1w images of 22 subjects at a 0.5 mm isotropic voxels size (see Krimmel et al. (2019) for details). These masks were used to replicate previous studies employing small region confound correction and enable comparability of results (Barrett et al., 2020; Stewart et al., 2024; Krimmel et al., 2019). For HCP7T data, the original CLA masks were used. At 3T, use of the original CLA masks led to scattered voxel extraction upon visual inspection. Therefore, the original CLA masks were dilated by 0.75 mm to guarantee the correct extraction of CLA voxels at the substantially larger functional voxel size.

For the correct definition of large-scale brain networks, ROI masks were manually created using functional activation maps derived from Neurosynth in conjunction with anatomical constraint masks derived from the Automatic Anatomical Labeling 3 (AAL3) atlas. Neurosynth is an online database for the large-scale, automated synthesis of fMRI data, allowing for the extraction of search-term-based functional activation maps based on 14.371 studies (Yarkoni et al., 2011). The AAL3 is an anatomical parcellation atlas based on a spatially normalized single-subject high-resolution T1w image provided by the Montreal Neurological Institute (MNI) (Rolls et al., 2020; Tzourio-Mazoyer et al., 2002). Search-term-based functional ROI masks were extracted from Neurosynth for the entire DMN, single nodes of the CEN and SN, and the thalamus, amygdala, and hippocampus as subcortical structures. Functional masks were thresholded and binarized using FSLeyes, only retaining peak activation clusters. Anatomical localization of the thresholded functional maps was performed using the AAL3 in MRIcron, after which the corresponding anatomical masks were extracted. Functional and anatomical masks were intersected, applying an anatomical constraint to functional activation maps, only retaining overlap of functional activation clusters with anatomical masks. Individual network component masks were extracted for the DMN, CEN, SN, and subcortical masks to retain bilateral and midline ROIs. Lastly, any overlap of different component masks was removed to ensure sound segregation of ROIs. These included bilateral overlap of the AG from the PPC, dilated CLA from the AI, and amygdala from the hippocampus, as well as midline overlap of the vACC from the mPFC, and left thalamus from the right thalamus. A summary of the Neurosynth search-terms and AAL3 constraints applied for the creation of each ROI mask, and a visualization of the corresponding ROI masks is provided in Table 1 and Fig. 2 of the extended data, respectively.

#### BOLD time-series extraction

The BOLD time-series for each ROI was extracted from the cleaned and preprocessed fMRI data using the same protocol across datasets. All functional voxels covered by the ROI masks were extracted from the T2*w functional BOLD sequence. To increase extraction precision in the PC3T dataset, extracted voxels for all cortical and subcortical ROIs were intersected with subject-specific gray matter masks obtained via the fMRIPrep preprocessing pipeline. Gray matter intersection of functional voxels was omitted for the dilated CLA masks, as only scattered voxels were retained when applied. No gray matter mask intersection was applied for HCP data. Finally, to unify the time-series of all voxels per ROI, principal component analysis (PCA) was applied, with the first principal component being used as a unified single time-series for each ROI for further analysis. A visualization of the extracted voxels for the bilateral CLA at 7T and 3T is provided in Figure 3 of the Extended Data.

#### Small region confound correction

Given the small anatomical size of the CLA and its proximity to the surrounding insula and putamen, accurate time-series extraction poses a challenge due to partial volume effects (Krimmel et al., 2019). To control potential CLA BOLD signal contamination by surrounding structures, small region confound correction (SRCC) was applied to the CLA time-series following the previously established protocol (Stewart et al., 2024; Barrett et al., 2020; Krimmel et al., 2019). CLA masks used for time-series extraction were dilated by 2 and 4 times the functional voxel size at 7T and 3T. For the HCP7T dataset, the original CLA masks were dilated by 3 and 6 mm, given the functional isotropic voxel size of 1.6 mm. For 3T datasets, the already dilated CLA masks were further dilated by 6 and 12 mm, given the functional isotropic voxel size of 3.2 mm in PC3T, which was not altered for HCP3T despite the difference in voxel size due to time constraints. These dilations were intersected to create shell masks following the shape of the CLA, which are two functional voxels apart from the CLA mask used for extraction. These shell masks were intersected with high-resolution mean insula and putamen masks to create flanking regions, overlapping with parts of the insula and putamen, while being distant enough to exclude any CLA BOLD signal being contained. Due to the difference in voxel size, the flanking regions covered more medial parts of the insula and lateral parts of the putamen, generally more proximal to the CLA, for the HCP7T dataset, whereas the inverse was the case for 3T data. Using these flanking masks, the time-series for the insula and putamen flanking regions were extracted and removed from the original CLA time-series via linear regression. The residual of the regression model was then used as the corrected CLA time-series, reducing signal contamination by the surrounding areas.

To evaluate the impact of SRCC on the BOLD timeseries, the explanatory power of each regression model was quantified using the coefficient of determination (R^2^). In this case, R^2^ measures the proportion of variance in the CLA time-series that is predictable from the surrounding flanking regions. For each subject, R^2^ was calculated using the residual sum of squares (unexplained variance) and the total sum of squares (total variance) for the left and right regression models. To evaluate the overall variance in the CLA time-series explained by the flanking regions, the mean and standard deviation of the R^2^ values from all subjects for the left and right regression models were calculated. To check for the success of SRCC, FC between the bilateral CLA to the ipsilateral putamen and insula was determined pre- and post-SRCC. FC was calculated using the Pearson correlation coefficient of functional voxels between the CLA and surrounding ROIs, with reduced FC reflecting a reduction in correlation between the BOLD time-series, interpreted as removal of partial volume effects. Lastly, we calculated the mean CLA BOLD variance across datasets, which was tested for a significant difference pre- and post-SRCC using a Mann-Whitney U test. For PC3T, we additionally distinguished between the baseline and psilocybin session, using BOLD variance as a proxy for regional activity, to test if psilocybin modulated CLA activity (Barrett et al., 2020).

#### Spectral dynamic causal modeling

Dynamic causal modeling (DCM) is a Bayesian modeling approach used to infer the effective connectivity (EC) between different brain regions based on indirect measures of neuronal activity, such as the fMRI BOLD time-series. Effective connectivity is defined as the influence one neuronal population exerts on another (Friston and Price, 2001). As such, it is distinct from functional connectivity (FC) as it estimates the directed causal influence between brain regions, as opposed to the statistical dependency between BOLD signals (Friston, 2011). EC is measured in Hertz (Hz), quantifying the rate of change in (neural states) per unit of time that one brain region causes in another. This, however, does not hold for the estimated self-connections, which, by default, are logarithmically transformed. This transformation enforces a negativity constraint, which is motivated by the biological plausibility of self-inhibition (Novelli et al., 2024). DCM comprises a biophysical forward model employing generative state-space modeling of stochastic neuronal dynamics and the hemodynamic response underlying the observed BOLD signal at the single-subject level ((Friston et al., 2003). The neuronal model is a linear stochastic differential equation that describes the dynamics of hidden neuronal states. It models how the activity in one brain region is influenced by the activity in other regions, as well as the endogenous fluctuations in neuronal activity. The observation function maps the modelled neuronal activity to the observed BOLD signal by modeling the biophysical processes involved in the haemodynamic response. As DCM is a Bayesian approach, all model parameters are equipped with prior distributions, which are updated based on the observed data to produce posterior distributions over the parameters. Using a Bayesian model inversion called Variational Laplace (Zeidman et al., 2023), these generative models are inverted to estimate the neuronal and hemodynamic parameters that expose the hidden neuronal state causing the BOLD signal. Spectral DCM (spDCM) constitutes a variation of DCM specifically developed for resting-state fMRI (rs-fMRI) without external stimulation. By fitting the model to the cross-spectral density of the measured BOLD signals, a second-order statistic that captures the correlations between time series at all time lags, spDCM can estimate the EC between brain regions, as well as the hemodynamic parameters and the spectrum of the endogenous fluctuations.

Effective connectivity was estimated using a custom shell and MATLAB scripts calling on functions of the DCM12 toolbox in SPM12. For each participant, four fully connected DCMs without exogenous input were fitted to the data for the CLA with the CEN, DMN, SN, and subcortical ROIs separately for each dataset. Each network model included four network components and the bilateral CLA, whereas the subcortical model included six bilateral subcortical components and the bilateral CLA. For all models, connectivity parameters for every possible ROI pair, as well as self-connections, were estimated, using default priors. For each dataset, the TR was adjusted in the DCM specification to match the TR used during imaging. The explained variance of all models exceeded an average of 80%, indicating successful and robust DCM model fit.

All reported EC estimates exceeded a posterior probability threshold of over 0.99, which is considered ‘very strong’ evidence.

#### Parametric empirical Bayes

Parametric empirical Bayes (PEB) was employed to combine DCM-based subject-level estimates at the group level. PEB employs a random effects model and Bayesian model reduction, estimating the group-level EC estimates while accounting for the varying degree of uncertainty in the estimates across subjects. For HCP data, PEB was employed to estimate the group-level EC, incorporating all participants and scans for each model. For PC3T data, PEB was employed to estimate group-level EC at baseline and its change under psilocybin. For this change analysis, the first regressor encodes the connectivity at baseline, and the second regressor encodes the difference between groups. Additionally, a PEB-based behavioral associations analysis was run separately only for the administration scan for each network using the MEQ30: Total and 11D-ASC composite scores as covariates. For this analysis, the first regressor in the design matrix represented the mean EC under psilocybin, and the second regressor the respective behavioral score as a covariate. To ensure interpretability of results, possible connections for associations were constrained to the ones for which a change in EC was identified between the baseline and psilcocybin scan in the initial analysis. All reported behavioral associations exceeded a posterior probability threshold of over 0.95, which is considered ‘strong’ evidence.

## Extended data

See attached file.

## Acknowledgements

The authors acknowledge the facilities and scientific and technical assistance of the National Imaging Facility (NIF), a National Collaborative Research Infrastructure Strategy (NCRIS) capability at Monash Biomedical Imaging (MBI), a Technology Research Platform at Monash University. A.R. is affiliated with The Wellcome Centre for Human Neuroimaging, supported by core funding from Wellcome [203147/Z/16/Z]. A.R. is a CIFAR Azrieli Global Scholar in the Brain, Mind & Consciousness We also acknowledge USONA Institute for providing psilocybin through their “Investigatational Drug Supply Program”.

## Data availability

The Human Connectome Project Young Adult (HCP-YA) 2017 release used in this study was publicly available upon license agreement at http://db.humanconnectome.org. Note that the new updated HCP-YA 2025 release will only be available at https://balsa.wustl.edu/project?project=HCP_YA. All PsiConnect data will be made fully open access through a OpenNeuro repository upon publication.

## Code availability

Open-source code for all data analysis pipelines will be made available on GitHub upon publication (https://github.com/razilab).

## Extended data

**Extended Data Fig. 1:**
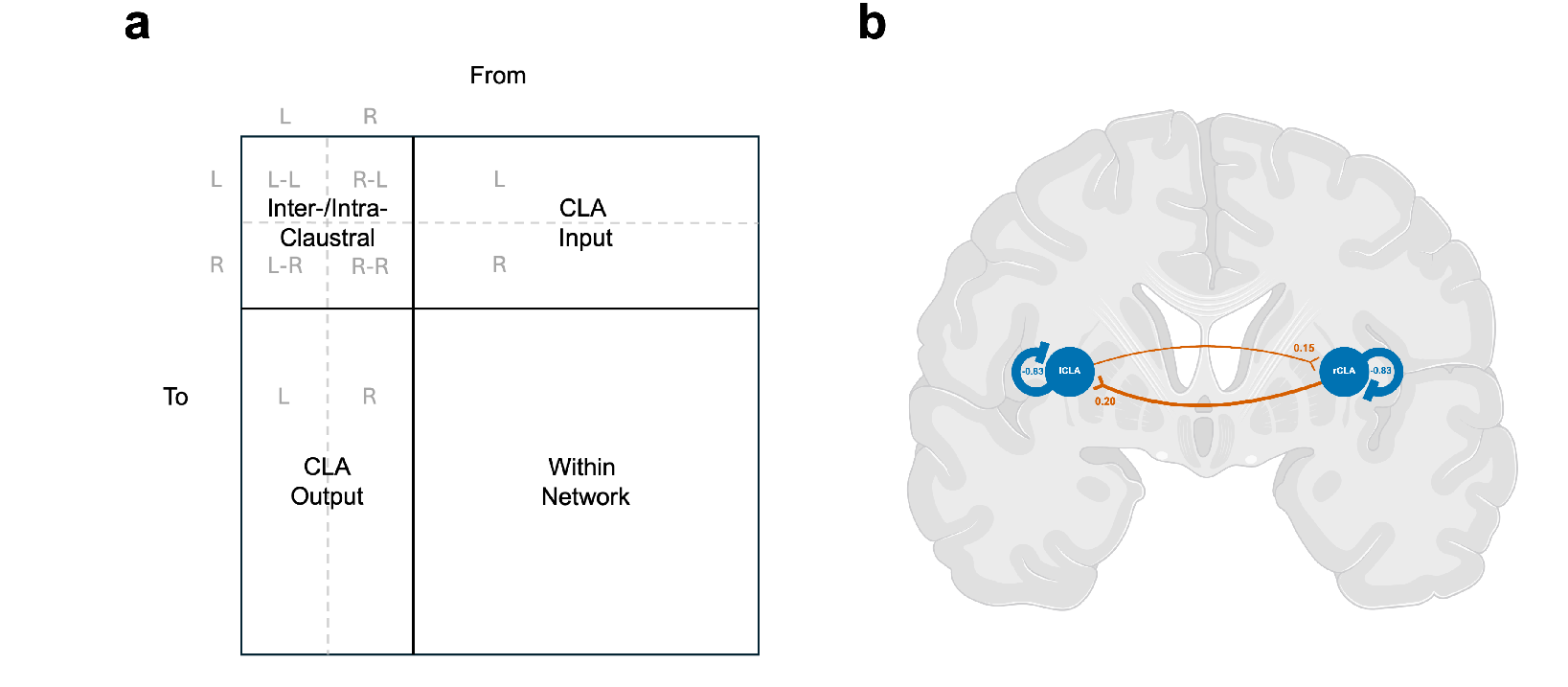
**a** Matrix layout to aid interpretability. **b** Mean EC estimates between CLA hemispheres and self-inhibition.

**Extended Data Fig. 2:**
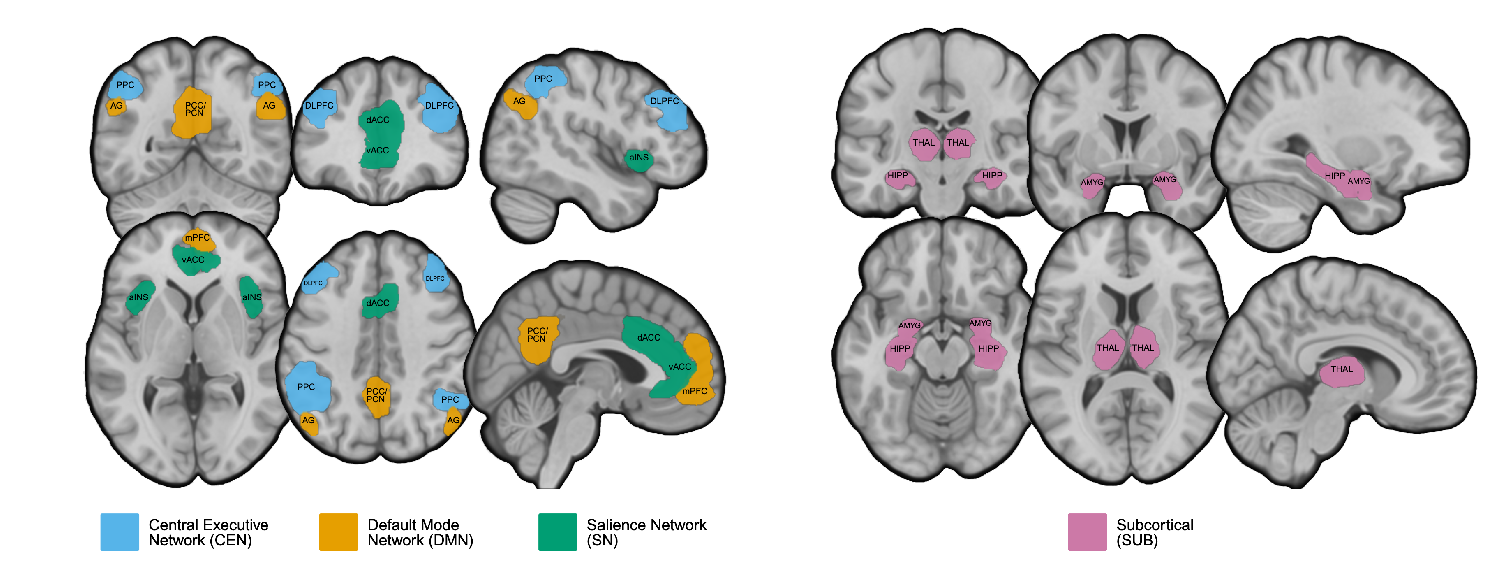
Regions of interest (ROIs) **A** Cortical network ROI masks **B** Subcortical ROI masks.

**Extended Data Fig. 3:**
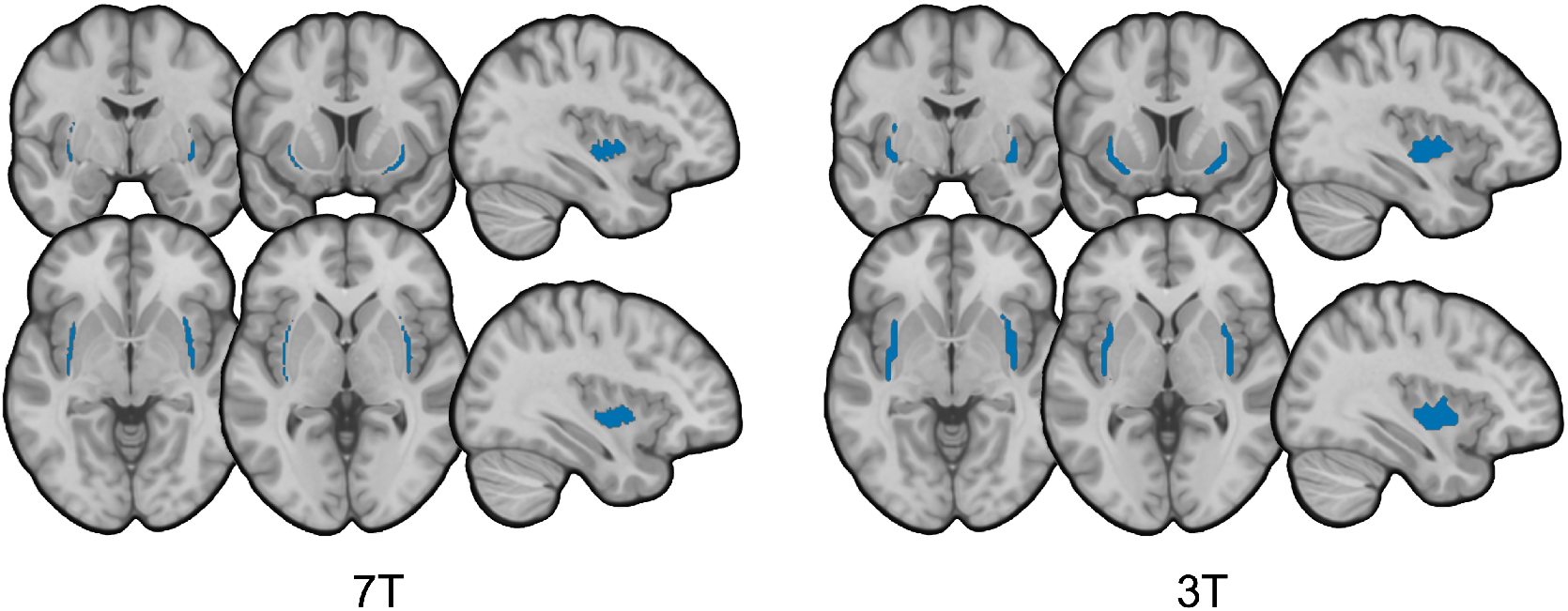
Extracted claustrum voxels

**Extended Data Table 1:**
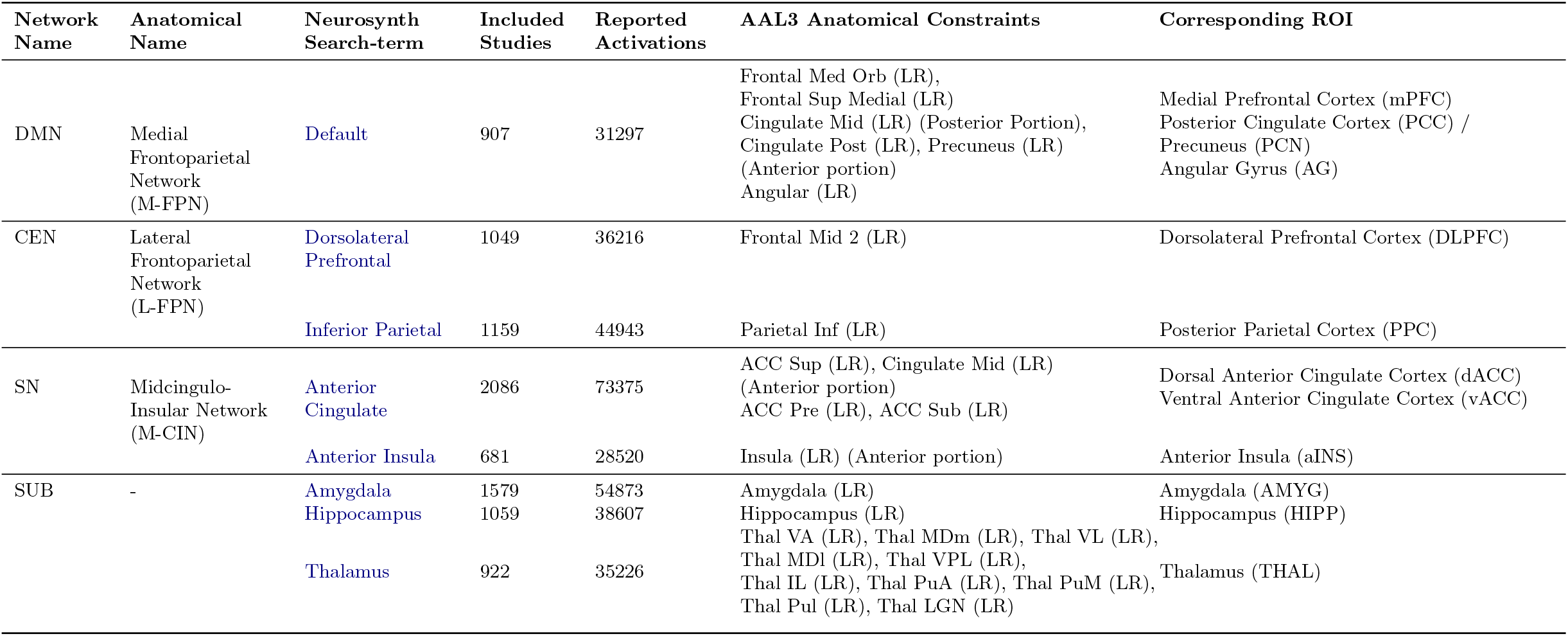
Neurosynth Functional Activation Search-terms and AAL3 Anantomical Mask Names

